# Structural characterization of NORAD reveals a stabilizing role of spacers and two new repeat units

**DOI:** 10.1101/2021.03.29.437547

**Authors:** Uciel Chorostecki, Ester Saus, Toni Gabaldón

## Abstract

Long non-coding RNAs (lncRNAs) can perform a variety of key cellular functions by interacting with proteins and other RNAs. Recent studies have shown that the function of IncRNAS are largely mediated by their structures. However, our structural knowledge for most IncRNAS is limited to sequence-based computational predictions. Non-coding RNA activated by DNA damage (NORAD) is an atypical lncRNA due to its abundant expression and high sequence conservation. NORAD regulates genomic stability by interacting with proteins and microRNAs. Previous sequence-based characterization has identified a modular organization of NORAD composed of several NORAD repeat units (NRUs). These units comprise the protein-binding elements and are separated by regular spacers of unknown function. Here, we experimentally determine for the first time the secondary structure of NORAD using the nextPARS approach. Our results suggest that the spacer regions provide structural stability to NRUs. Furthermore, we uncover two previously-unreported NRUs, and determine the core structural motifs conserved across NRUs. Overall, these findings will help to elucidate the function and evolution of NORAD.

## Introduction

Long non-coding RNAs (lncRNAs) constitute a heterogeneous family of RNA transcripts defined by their length (> 200 nucleotides (nt)), and their lack of protein-coding potential. Most lncRNAs are expressed at low levels and in a tissue- or cell type-specific manner, and are poorly conserved both at the sequence and exon-intron structure levels [1,2]. Over the last years, an increasing number of lncRNAs have been associated with a variety of cellular and developmental functions [3–5], and have been shown to be altered in various human diseases [6]. However, the functions and structures of the vast majority of lncRNAs, remain poorly understood.

LncRNAs can form complex secondary structures, which tend to be more conserved than primary sequences and are thought to mediate their biological functions [7,8]. However, controversy remains as to whether secondary structures of lncRNAs are evolutionarily conserved [9]. Although molecular probing techniques allow assessing RNA structures folded *in vitro* the large size of many lncRNAs poses a significant challenge. As a result, only a few IncRNA structures have been experimentally characterized *in vitro* [9], including those of HOTAIR [10], RepA [11], Xist [12,13], ncSRA [14] and lincRNAp21 [14,15]).

Many sequences, including riboswitches, mRNAs and long non-coding RNAs [9], are expected to fold into a variety of competing structures that can coexist. To account for this structural diversity, several methods have been recently developed that allow to study alternative RNA conformations using experimental data [16–20].

The non-coding RNA activated by DNA damage (NORAD) is a 5.3 kilobases (kb) long IncRNA that is widely expressed and conserved across mammals. Previous sequence analyses of this IncRNA have described the presence of several repeated elements - NORAD repeat units or NRUs - that are predicted to fold into similar secondary structures and tend to be conserved [21,22]. NORAD maintains genomic stability by sequestering RNA-binding PUMILIO proteins (PUM1 and PUM2), thereby preventing them from interacting with their targets [21]. PUMILIO proteins act by repressing the translation of specific target mRNAs by binding to PUMILIO response elements (PRE) present in their 3’ untranslated region (3’UTR) [23]; [23,24]. In the absence of NORAD, PUMILIO proteins drive chromosomal instability caused by DNA repair, hyperactively repressing mitotic, and DNA replication factors [21].

In addition, NORAD acts as a competitive endogenous RNA (ceRNA) and regulates cancer progression by sponging MicroRNAs (miRNAs) [25,26]. MiRNAs are small regulatory RNAs of about ~20 nt in length that play critical regulatory roles in eukaryotic cells where they mediate post-transcriptional silencing of specific genes [27–29].

Despite the increasing relevance and known functions of NORAD, the empirical characterization of its structure is still lacking. To shed light on the function and evolution of NORAD, we here characterized its secondary structure using experimental probing with the nextPARS approach [30]. In addition, we assessed how changes in sequence context and shifts in temperature impact the structure of NRUs and other relevant NORAD motifs. Our results provide evidence that the spacer sequences provide structural stability to NRUs and that the structure around conserved SAM68-binding sites is more robust to thermal shifts than other positions in the NORAD sequence. Finally, we define the core structural motifs conserved across NRUs, and uncover the presence of two previously-unreported NRUs.

## Materials and Methods

### Sample preparation

The structure of several NORAD fragments was experimentally probed using nextPARS (PMID: 29358234) at three different temperatures (23 °C, 37 °C and 55 °C) by spiking the fragments of interest into fungal total or polyA+ RNA samples.

The probed NORAD fragments were the following: three RNA fragments covering the full-length NORAD lncRNA (called NORAD#1, NORAD#2 and NORAD#3, overlapping 1903, 1862 and 1614 bases, respectively), and 11 NORAD repetitive units previously described [22] with their surrounding regions (NRU#1, NRU#2, NRU#3, NRU#4, NRU#6 NRU#7, NRU#8, NRU#9, NRU#10, NRU#11 and NRU#12, overlapping 179 nt each) [Supplementary table S1]. To produce these fragments, NORAD was cloned into the pUC57 vector by GenScript Biotech (Netherlands). The plasmid was transformed in DH5α *E. coli* competent cells previously prepared in our lab according to manufacturer’s instructions (Mix & Go E. coli Transformation Kit & Buffer Set, ZymoResearch). Single colonies were grown in LB+Ampicillin medium (37 °C, 220 rpm, overnight), and plasmids were purified using QIAprep Spin Miniprep kit according to the manufacturer’s instructions (Qiagen). PCRs were performed to amplify and linearize the different NORAD fragments using the Pfu mix (#2021, DongSheng Biotech) and a 0.4 μM final concentration of each primer (forward and reverse) in a total volume of 40 μl. Supplementary Table S1 shows primer sequences and amplicon sizes and Supplementary Table S2 specifies PCR conditions used per each fragment. PCR amplicons were purified using a QIAquick PCR purification kit according to the manufacturer’s instructions (Qiagen) and then they were Sanger sequenced, to confirm that no mutation had been introduced in the fragments of interest. Then, all NORAD fragments already linearized were transcribed *in vitro* using the T7 RiboMax Large-scale RNA production system according to the manufacturer’s instructions (Promega). Finally, RNAs of interest were size-selected and purified using Novex-TBE Urea gels according to the manufacturer’s instructions (Life Technologies). Final quality control of the purified RNAs was performed using the Agilent 2100 Bioanalyzer with the RNA 6000 Pico LabChip Kit (Agilent) and the Qubit Fluorometer with the Qubit RNA BR (Broad-Range) Assay Kit (ThermoFisher Scientific). Ten other RNA molecules were spiked into the samples (TETp4p6, TETp9-9.1, hSRA, mSRA, ROX2, GAS5, HOTAIR_NCBIBI_RINN, 372_BRAVEHEART, 373_BRAVEHEART, B2 and U1) [Supplementary table S3]

### Secondary structure probing with NextPARS

For the enzymatic probing of RNA samples at different temperatures (23 °C, 37 °C and 55 °C), we used the nextPARS protocol [30], adapting it when necessary for the probing at higher temperatures. Briefly, 2 μg of PolyA+ RNA or total RNA were mixed with 20 ng of each NORAD RNA fragment and were brought to a final volume of 80 μl with nuclease-free water. Samples were first denatured at 90 °C for 2 min and immediately cooled down to 4 °C on ice for 2 min. Then, 10 μl of ice-cold 10X RNA structure buffer (Ambion) were added to the samples and mixed by pipetting up and down several times. When probing at 23 °C, RNA samples were subsequently brought from 4 °C to 23 °C, in about 20 min (1 °C per min). For samples probed at 37 °C and 55 °C, the temperature was brought from 4 °C to 30 °C (1 min per each °C), samples were incubated 5 min at 30 °C, and then brought to 37 °C and 55 °C, respectively, for enzymatic digestion. Finally, 10 μl of the corresponding dilutions of RNase V1 (Ambion), S1 nuclease (Fermentas), or nuclease-free water were added to the V1-digested, S1-digested and undigested samples, respectively. The enzyme concentration used was different depending on the probing temperature, to compensate for the effect of higher temperatures to prevent over-digestion of the RNA samples and according to previous studies [31]. Thus, the final reaction contained 0.03 U, 0.015 U or 0.0075 U of RNAse V1, and 200 U, 100 U or 50 U of S1 nuclease for samples probed at 23 °C, 37 °C or 55 °C, respectively. After mixing by pipetting, samples were incubated at each corresponding temperature for 15 minutes more. The RNAs were purified using RNeasy MiniElute Cleanup kit following the manufacturer’s instructions (Qiagen). Quality controls with NanoDrop 1000 Spectrophotometer (Thermo Scientific) and Agilent 2100 Bioanalyzer with the RNA 6000 Pico LabChip® Kit (Agilent) were performed before and after the digestion, to confirm that samples were not digested with the initial denaturation or over-digested after the enzymatic treatment.

### Library preparation and sequencing

We used the TruSeq Small RNA Sample Preparation Kit (Illumina) to prepare the libraries for further sequencing. We initially performed a phosphatase treatment incubating 16 μl of the digested samples for 30 min at 37 °C and 5 min at 65 °C mixed with 2.5 μl of 10X phosphatase buffer, 2.5 μl of nuclease-free water, 1 μl of RNAse inhibitor and 3 μl of Antarctic phosphatase (New England BioLabs Inc.). We then performed a kinase treatment adding 4 μl of T4 Polynucleotide Kinase (PNK, New England BioLabs Inc.), 5 μl of 10X PNK buffer, 10 μl of ATP 10 mM, 1 μl of RNAse inhibitor and nuclease-free water up to a total volume of 50 μl, and incubating the samples 1 hour at 37 °C. Samples were then purified using RNeasy MiniElute Cleanup kit following manufacturer’s instructions (Qiagen) with a 10 μl RNase-free water final elution step. Samples were concentrated using a centrifugal evaporator Speed Vac® to a final volume of 5 μl and we started the TruSeq Small RNA Sample Preparation Kit (Illumina) protocol according to manufacturer’s instructions. After the final purification of the cDNA libraries, we performed quality control of each library using Agilent 2100 bioanalyzer with the DNA 1000 Kit (Agilent). Libraries were sequenced in single-reads with read lengths of 50 nucleotides in Illumina HiSeq2500 sequencers at the Genomics Unit of the CRG.

### Computation of the nextPARS scores

The computation of the nextPARS scores was obtained following the protocol in [30,32]. Briefly, we removed adapters with cutadapt and mapped the reads using STAR [33]. Then, we used in-house scripts for parsing the mapped reads and retrieved the 5’ -end position in the NORAD transcript. This results in a digestion profile that indicates the number of cuts per position of the transcript. To obtain the scores from nextPARS probing experiments we used as an input the digestion profile. The output contains a structural profile of the NORAD transcript (single-or double-stranded state), with a score for each nucleotide that ranges from −1.0 (highest preference for single strand) to 1.0 (highest preference for double-strand) (https://github.com/Gabaldonlab/nextPARS_docker).

### RNA secondary structure prediction and visualization

To obtain the secondary structure of NORAD and NRUs we use RNAstructure software [34]. Using nextPARS2SHAPE v1.0 script (https://github.com/Gabaldonlab/MutiFolds/) we converted nextPARS score to SHAPE-like normalized reactivities that were used to provide pseudoenergy restraints to the Fold software. The consensus structure models have been generated using the R2R software [35]. RNA structures were constructed using VARNA (Version 3-93) [36].

### Identification of repetitive units in NORAD

To identify new repetitive elements in NORAD we first performed a multiple sequence alignment with T-Coffee (Version_11.00.8cbe486) [37] using the regions of the 12 described NRUs. Then, a covariance model (CM) was built using Infernal 1.1.3. [38] cmbuild on the 12 NRUs sequences with --noss parameter to ignore the secondary structure annotation. Then, we calibrated the CM with cmcalibrate. Next, cmsearch was used to search for the CM against NORAD and the 14 sequences below the E-value cutoff (0.01) were retrieved.

### Prediction of multiple folding conformations, guided by nextPARS data

Rsample (Version 6.2) [19] was used to calculate the partition function for the NRUs from NORAD sequence to determine the multiple conformations. To perform a complete Rsample calculation, we performed the following steps. First, we converted the nextPARS score to SHAPE-like reactivities and then we ran Rsample using the SHAPE-like constraint to produce a Partition Save File (PFS). After that, we ran stochastic, using this PFS file as input, and we created a CT file with Boltzmann ensemble of 1,000 structures. Finally using the R script from Rsample (RsampleCluster.R) we use the CT file to calculate the optimal number of clusters and their centroids to model the NRUs sequences that populate multiple structures, including the relative probabilities of those structures. To evaluate conserved structural elements within the ensemble of structures of the NRUs, we used the Beagle (BEar Alignment Global and Local) software [39]. We performed an all-vs-all alignment for all the NRUs to identify common secondary structures and compared them with the most stable structure.

### Statistical methods

Statistics were performed using Wilcoxon rank-sum and Kruskal-Wallis tests. The similarity of nextPARS scores between the same residues in NRU segments folded independently or within each piece and larger NORAD fragments was measured by Pearson’s correlation. The Gardner-Altman estimation plots were produced using the python version of dabest [40].

### Deposited data

Raw sequencing data of nextPARS experiments were deposited in the Sequence Read Archive under the BioProject ID PRJNA714002. (Reviewer’s access https://dataview.ncbi.nlm.nih.gov/object/PRJNA714002?reviewer=c05bml4s9go209k1vvhhs4pes7).

## Results

### Structural characterization of NORAD

Resolving the structure of NORAD *in vitro* is exceptionally challenging because of the large size of this transcript. To circumvent this problem, we synthesized, cloned, and *in vitro* transcribed (see Materials and Methods) three overlapping NORAD fragments [Fig. 1A], and subsequently subjected them to enzymatic probing using the nextPARS method [30]. This approach renders single-nucleotide resolution measurements of *in vitro* secondary structure, which can be represented by a structural profile (single- or double-stranded state), providing a score for each residue, ranging from −1.0 (highest preference for single strand) to 1.0 (highest preference for double-strand). We converted the nextPARS score to SHAPE-like reactivity and used this to provide constraints for secondary structure prediction [Fig. 1B]. Our results show that most of the NRUs form independently-folded structures (NRU#2, NRU#3, NRU#5, NRU#7, NRU#8, NRU#9, NRU#11 and NRU#12) but some of them seem to interact with each other or with other NORAD regions [Fig. 1B].

**Figure 1.**
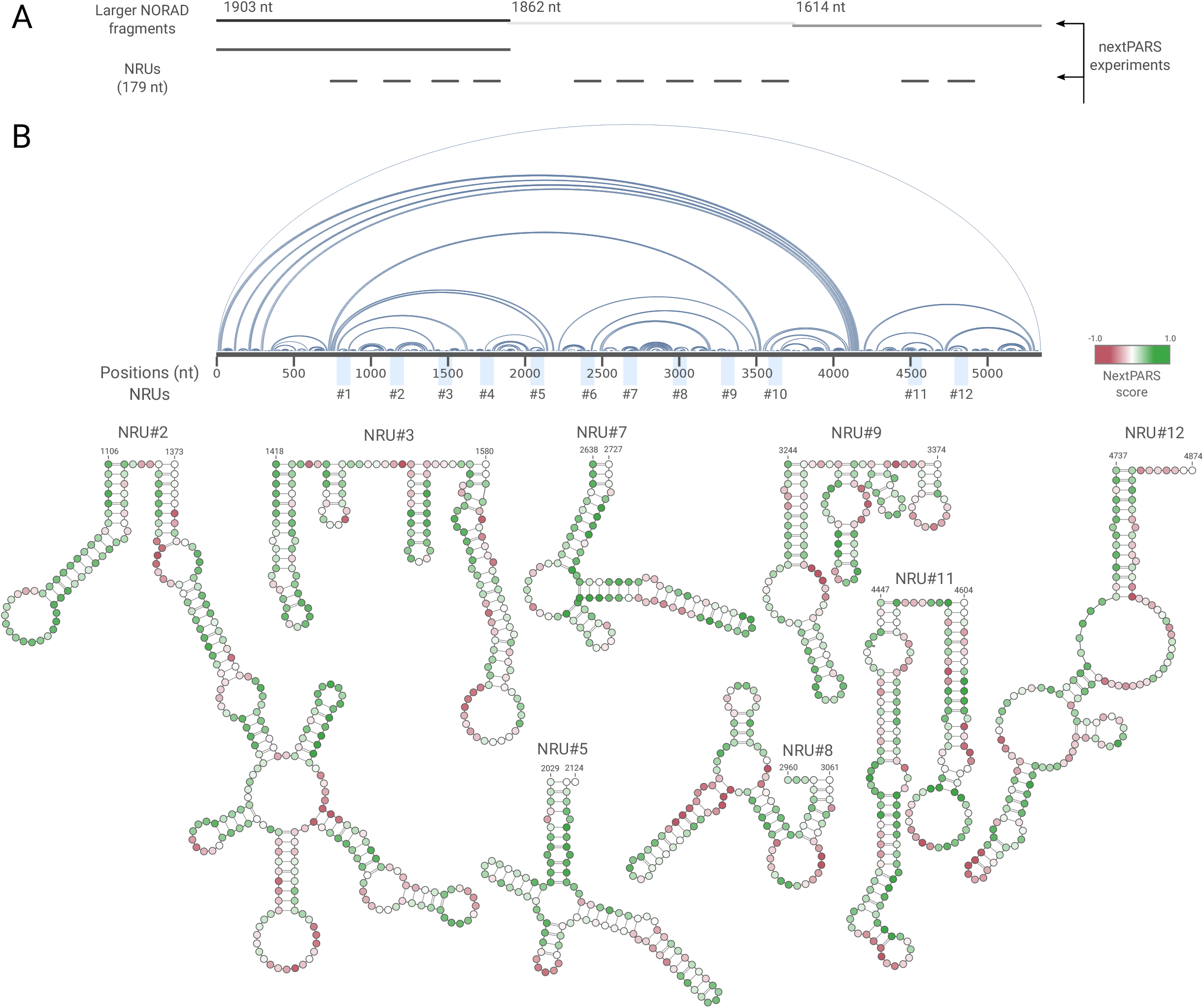
The landscape of NRUs structures using experimental nextPARS data. **A.** Overview of the nextPARS experiments. NORAD fragments were probed in three RNA fragments covering the full-length NORAD lncRNA (overlapping 1903, 1862 and 1614 bases, respectively), and 11 NRUs with their surrounding regions (overlapping 179 nt each). **B.** On the top, an arc plot depicting the interactions of the secondary structure inferred from the nextPARS profiles using RNAstructure software. The core sequences of the NRUs (89nt) are marked alongside NORAD. On the bottom, the secondary structure of NRUs derived from the full-length NORAD nextPARS experiment. Scores from full-length NORAD nextPARS profiles are depicted by colored nucleotides.

To analyze NRUs structures in detail, and to assess the effect of the NORAD sequence context in NRU folding, we designed non-overlapping 179 nt long fragments covering 11 of the 12 previously described NRUs [22] and performed nextPARS experiments [Fig. 1A]. To identify structural subdomains in NORAD, we followed a strategy previously used with HOTAIR [10,41]. In this approach, the full-length lncRNA profile is compared to the profiles of individual RNA segments to identify potential independently-folded sub-domains. Comparing the nextPARS profile of the NRUs with the same sequence in the context of the full NORAD molecule, shows a high correlation, with a p-value < 0.01 [Supplementary Figure S1]. Pearson’s correlation coefficient values (*ρ*) are expected to be higher for fragments that do not break elements of the secondary structure observed in the full-length molecule. This was the case for the eleven NRUs analyzed (*ρ* > 0.8). Correlations values were particularly high (*ρ* > 0.9) for some of the NRUs (NRU#1, NRU#4, NRU#8, NRU#10 and NRU#12), suggesting that these regions form more stable local structures [Supplementary Figure S1].

### Spacer sequences provide structural stability to NRUs

To evaluate the robustness and stability of NORAD secondary structure, we performed nextPARS experiments at three different temperatures: 23 °C, 37 °C and 55 °C to study the alterations prompted by these temperature shifts as a means of structural perturbation [Fig. 2A]. We compared nextPARS scores obtained in equivalent residues when probed in NRU-specific fragments or in larger NORAD fragments, and when probed at different temperatures in the context of the two molecular sizes. On average, the correlation between secondary structures was higher across varying temperatures than when NRUs structures were compared between those folded in isolated fragments with larger NORAD fragments [Fig. 2A,B]. These results suggest that the molecular context provided by the surrounding NORAD spacer sequences has a larger influence on the structure of NRU than sharp shifts in temperature.

**Figure 2.**
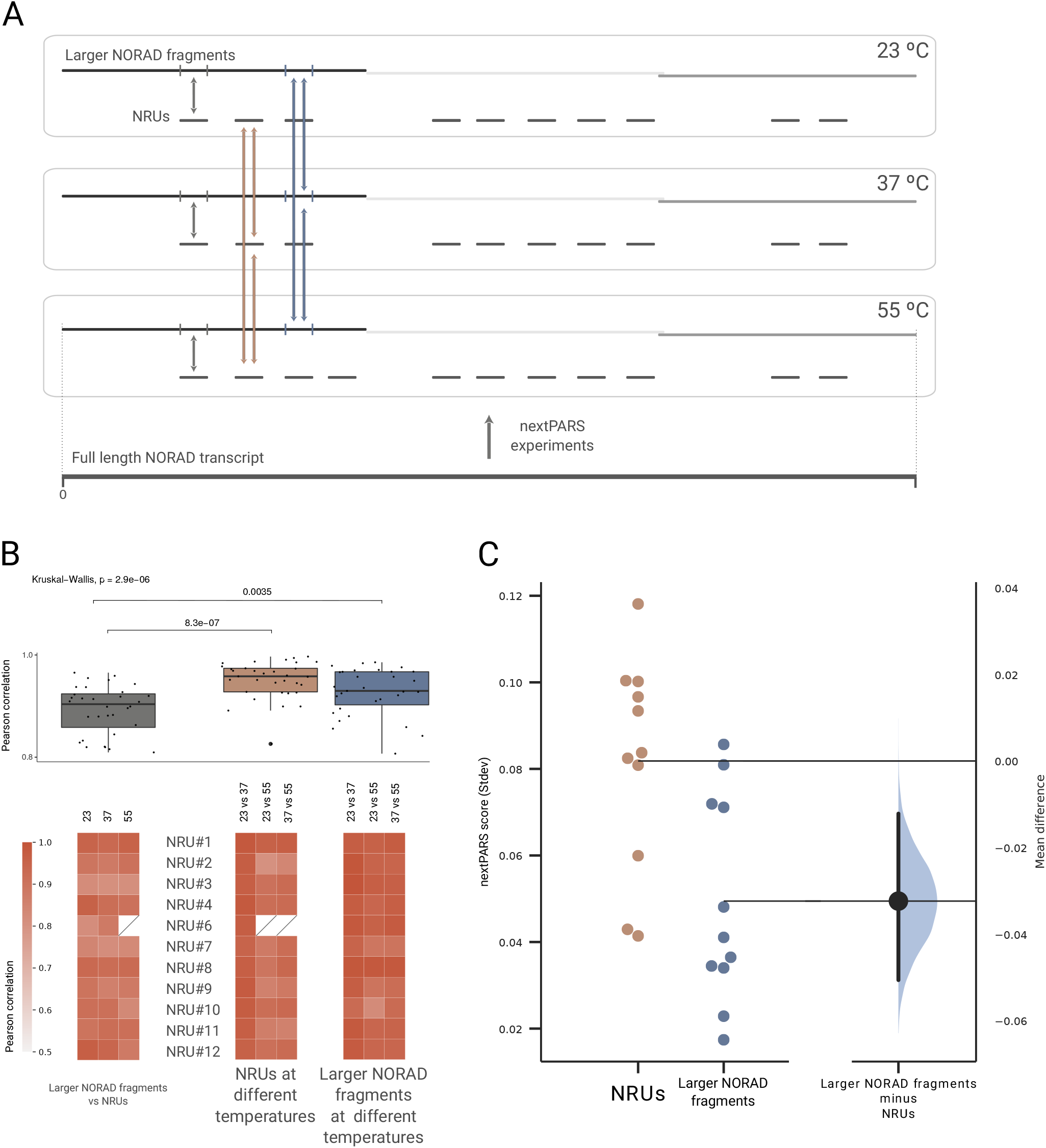
Molecular context provides structural stability to NRUs. **A.** Overview of nextPARS experiments at three different temperatures: 23 °C, 37 °C and 55 °C. Arrows showing different comparisons: in grey NRUs compared between those folded in isolated fragments with larger NORAD fragments, in brown comparison of each NRUs at different temperatures by pairs, and in blue the comparison of NRUs at different temperatures by pairs, in the context of the larger NORAD fragments experiments. **B.** On the top, a box plot of Pearson correlations of the three different comparisons described in A, with individual observations on top of boxes. On the bottom, a heatmap showing the Pearson correlation coefficients for each of the comparisons. **C.** Gardner-Altman estimation plot showing the mean difference between the average of the standard deviation of nextPARS score at the three different temperatures for the NRUs and for the larger NORAD fragments. Both groups are plotted on the left axes; the mean difference is plotted on a floating axis on the right as a bootstrap sampling distribution. The mean difference is depicted as a dot; the 95% confidence interval is indicated by the ends of the vertical error bar.

Then, we measured the stability of NRUs by averaging the standard deviation of nextPARS scores at different temperatures [Fig. 2C]. Importantly, we observed that structural variability across temperatures was higher in NRUs fragments compared to larger NORAD fragments, supporting a stabilizing role of the molecular context when NRUs are embedded within larger NORAD fragments [Fig. 2C]. To ensure that the observed structural variability differences are not due to the effect of sequence length (i.e. longer molecules having higher stabilities), we use the information of nextPARS experiments of ten RNA molecules that were spiked with NORAD samples, with variable size [30], and measured the structural stability at different temperatures. Our results did not detect any statistically significant correlation between stability and sequence length [Supplementary Figure S2], suggesting our results cannot simply be attributed to differences in length between NRU-specific, and larger NORAD fragments.

### Conserved SAM68-binding sites in NORAD are particularly robust to structural perturbation by temperature shift

SAM68 (KHDRBS1) is an RNA-binding protein (RBP) that performs several functions in the nucleus and cytoplasm [42]. It is required for efficient recruitment of PUMILIO2 protein (PUM2) to NORAD and the regulation of Pum activity by NORAD [43–45]. SAM68 recognizes UAAA motifs [46], which in NORAD are present in 16 motif pairs separated by linkers of 15-35 bases. These motifs show higher evolutionary conservation as compared to the rest of the NORAD sequence [43]. When comparing nextPARS score variability at different temperatures for NRU-specific fragments in positions of UAAA motifs to other positions of NRUs, we observed that the motif sites were less variable as compared to the remaining residues in the NRUs (p-value = 0.010148) [Fig. 3A]. To further investigate the association between sequence conservation and structural flexibility in UAAA motifs, we measured the correlation between the evolutionary conservation as estimated by the PhyloP score (based on the 100-way vertebrate alignment) and the variability of structural profiles in the position of all UAAA motifs. First, we measured the structural variability of the UAAA motifs using the nextPARS probing data from NRU-specific fragments and found no significant correlation between the level of conservation and the amount of structural variability [Fig. 3B]. Then, we calculated the same measure of correlation but using the nextPARS data from the larger NORAD fragments and observed a significant negative correlation between sequence conservation and structural variability within UAAA sites- the higher the sequence conservation, the lower the structural variability at these positions - [Fig. 3B]. We obtained similar results using PhastCons score or Phylop score at different conservation levels [Fig. 3B]. These results suggest that SAM68-binding sites (UAAA motif) are robust to structural perturbation by temperature shift.

**Figure 3.**
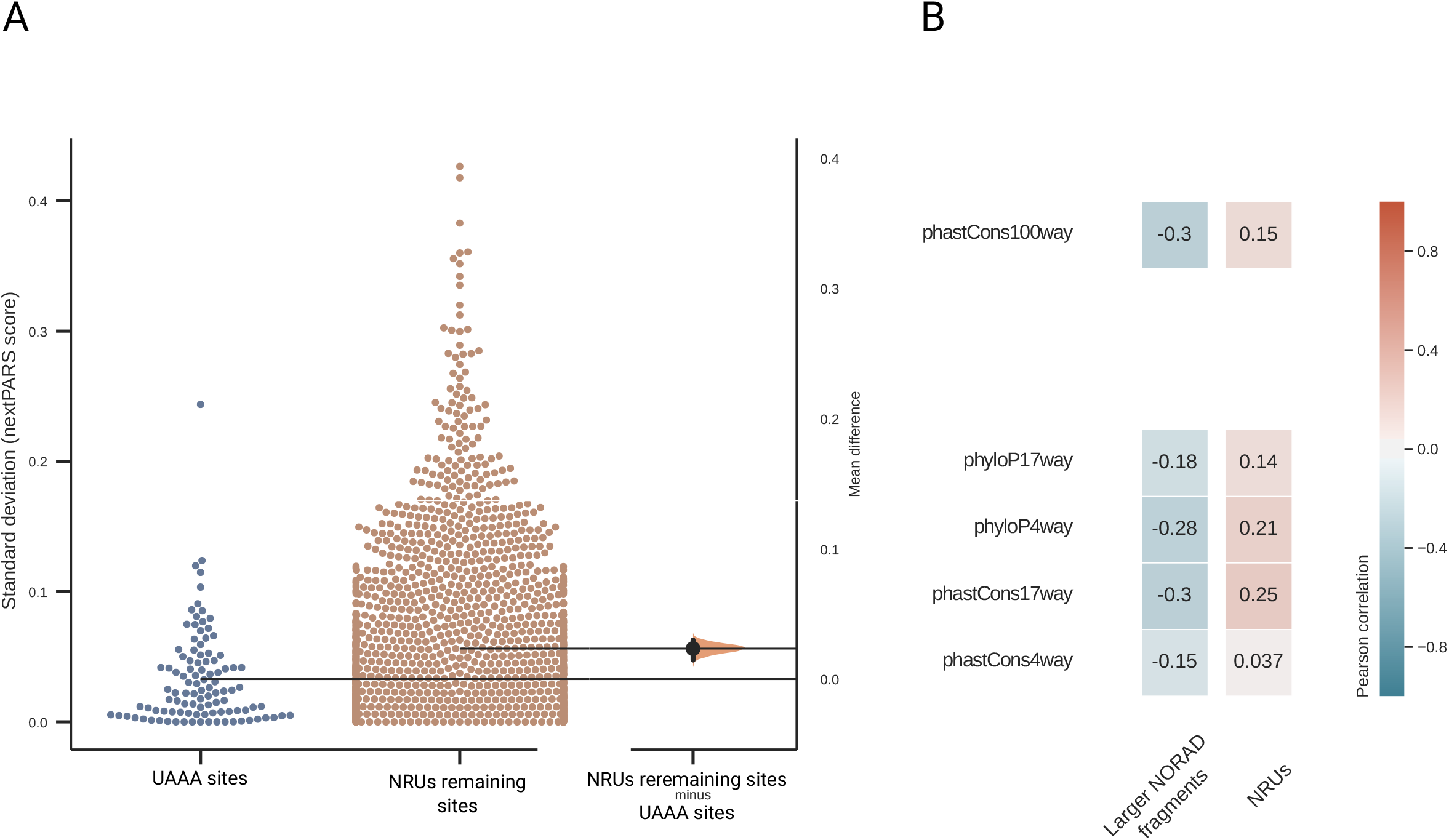
Comparison of positions of SAM68-binding sites against the remaining NRUs positions. **A.** Gardner-Altman estimation plot showing the mean difference between the standard deviation of nextPARS score at the three different temperatures for SAM68-binding sites positions and the NRUs remaining positions. Both groups are plotted on the left axes; the mean difference is plotted on a floating axis on the right as a bootstrap sampling distribution. The mean difference is depicted as a dot; the 95% confidence interval is indicated by the ends of the vertical error bar. **B.** A heatmap showing the Pearson correlations coefficients for nextPARS score variability at different temperatures in positions of UAAA motifs to other positions of NRUs. We show different correlations for PhastCons score or Phylop score at different conservation levels, and using data from isolated NRUs and NRUs in the context of larger NORAD fragments.

### Considering structural diversity of NRUs to study functional and evolutionary relationships

We next set out to investigate other NORAD conserved elements beyond the known protein binding sites. We used our empirical data to study the two previously predicted hairpin structures present in the NRUs core, as such conserved secondary structure motifs are rarely detectable in lncRNAs [22]. We produced a multiple sequence alignment to derive a consensus secondary structure from the eight NRUs displaying a clear conserved long stem-loop hairpin [43] (NRU#1, NRU#3-8 and NRU#10). In the predicted consensus structure [Fig. 4A], we observed a short stem-loop structure comprising four paired bases and a variable loop followed by a second large stem-loop structure with nine base pairs.

**Figure 4.**
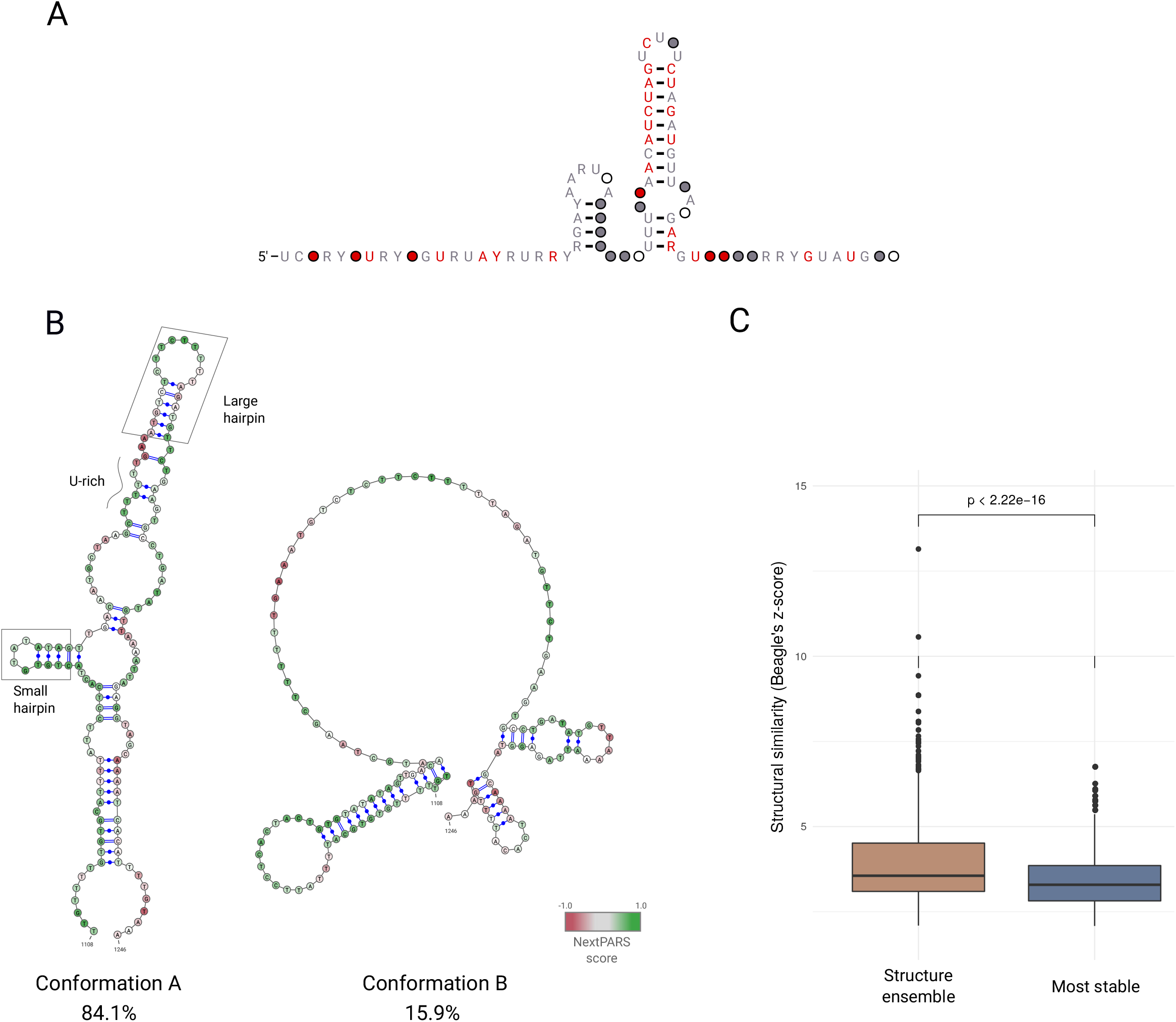
Comparison of alternative structures of NRUs and obtained consensus structure. **A.** The consensus secondary structure for the core region of NRUs, derived by using a multiple sequence alignment to derive a from the eight NRUs that have a clear conserved long stem-loop hairpin (NRU#1, NRU#3-8 and NRU#10). **B.** Secondary structure conformations A and B, from NRU#2 obtained using Rsample software in combination with nextPARS from larger NORAD fragments experiments. Scores from nextPARS scores are depicted by colored nucleotides. Known structural and sequence motifs are marked in the conformation A. **C.** A box plot showing the structural similarity (Beagle’s z-score) elements found by Beagle software, for the structure ensemble alignment and for the most stable structure alignment.

Many sequences including mRNA, lncRNAs and riboswitches are expected to fold in multiple, coexisting structures [9]. To investigate this in the case of NORAD, we used Rsample [19] in combination with nextPARS data to predict the presence of alternative RNA structures for NRUs, and studied these conformations. NRU#2 was one of the four NRUs that was excluded from the analysis of the consensus structure, as this unit did not contain most of the repeat motifs in comparison to the rest of the NRUs [22]. When we predicted the structure of NRU#2, we observed that it was folded in two alternative conformations. The major conformation (inferred to represent 84.1% of the molecules in the mix) was predicted to form a large stem-loop after a U-rich stretch, and a small stem-loop before that U-rich [Fig. 4B]. These detected structure motifs were similar to the ones detected in some of the characterized NRUs (NRU#1, NRU#3-8 and NRU#10) [22]. In contrast, in the minor conformation (15.9%) or in the computationally predicted conformation, we could not detect the combination of sequence and structure motifs present in the other NRUs that were previously characterized [Fig. 4B and Supplementary Figure S3A]. This indicates that considering the possibility of competing structural conformations enhances our ability to detect structural similarity that is otherwise missed by standard approaches based on a single stable structure.

Then, we searched for shared structurally related regions among the different NRUs, considering alternative conformations. For that purpose, we used the Beagle software [39] to perform an all versus all comparison of the most stable structure in all NRUs fragments. Then, we did the same all versus all comparisons but using the complete ensemble of alternative structures. We observed that the structural similarity (z-score) was higher for the ensemble of structures, as compared with the most stable one (*p*-value = 0.00095) [Fig. 4C], meaning again that using the alternative structures that may coexist within NORAD could be more accurate than using only the most stable one.

### Discovery of two previously uncharacterized NRUs

Previous comparison of human NORAD sequences revealed that this lncRNA could be decomposed into at least 12 repeating units of ~300 nt each [43]. When we studied the relative positions of the core sequence of the NRUs throughout NORAD, we noticed that consecutive units were separated by a sequence spacers of similar length (between ~270 nt and ~340 nt), with the exception of NRU#10 and NRU#11, which were separated by ~900 nt [Fig. 5A]. To investigate the presence of novel NRUs, we used infernal [38] to derive a model for NRUs (see Materials and Methods). Using this model on NORAD, we identified the 12 known NRUs, as expected, but we also detected the presence of two additional NRUs that we termed NRU#10a and NRU#10b. Interestingly, these units appear within the unusually long intervening region between NRU#10 and NRU#11 and are separated by the approximately same distance as the rest of the NRUs [Fig. 5A]. Multiple sequence alignment of the core sequences of the known and newly identified NRUs showed a high degree of sequence similarity [Supplementary figure S4A]. Furthermore, we noticed a common sequence motif located at ~50 nt downstream of the end of the core of NRU#10a and other NRUs. [Supplementary figure S4B]. Moreover, in the new NRUs, we could identify most of the sequence and structure motifs typical for known NRUs, including the presence of SAM68-binding sites within the NRU or nearby [Fig. 5B] [43].

**Figure 5.**
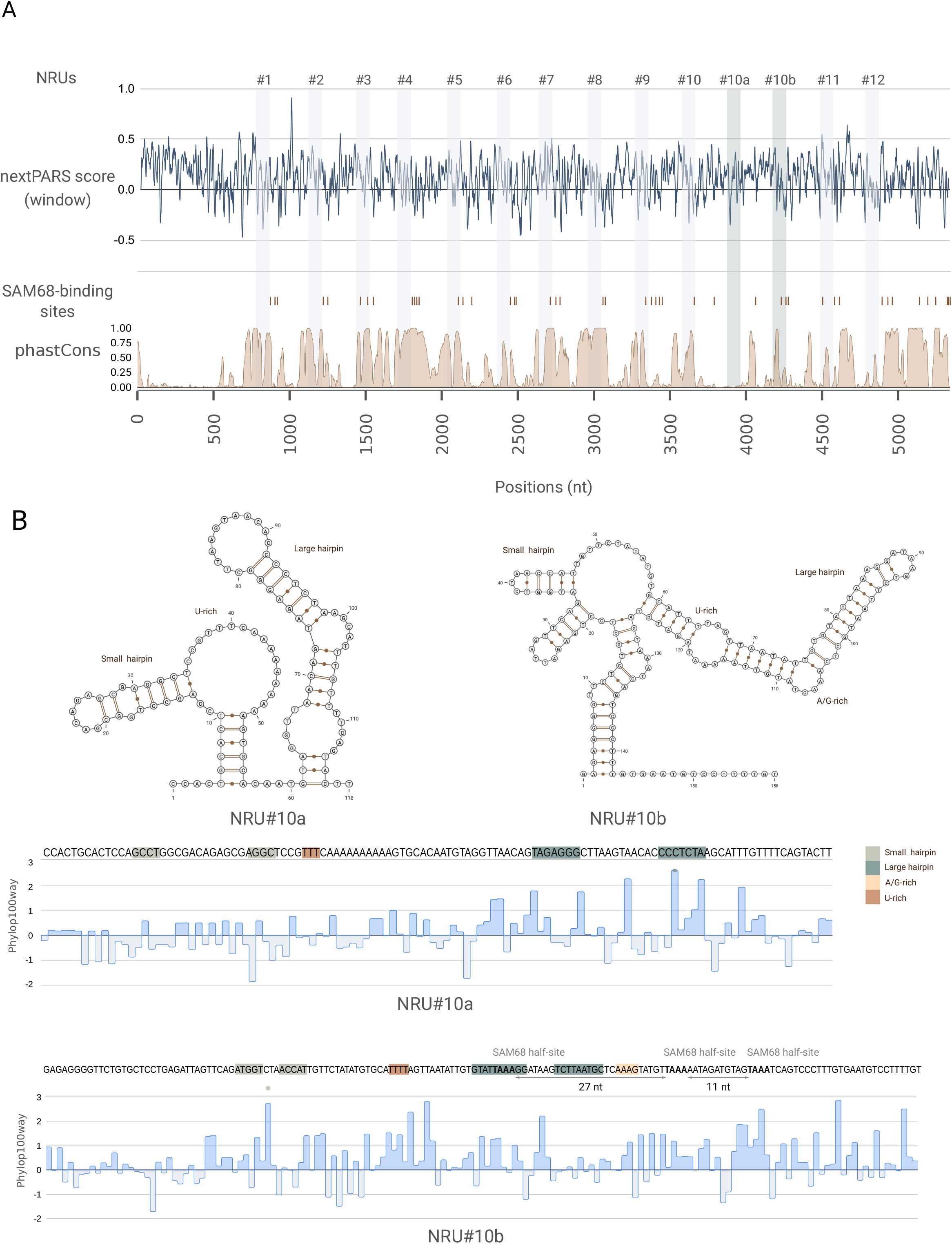
Discovery of two previously uncharacterized NRUs. **A.** Overview of the relative positions of the core sequence of the NRUs along with NORAD. The core positions of previously characterized NRUs are marked with grey bars, and the two new NRUs (NRU#10a and NRU#10b) are marked with green bars. The nextPARS score derived from the larger NORAD fragments experiment is shown, by an average window of 10nt. SAM68-binding sites are marked with brown lines. PhastCons score (based on the 100-way vertebrate alignment) is shown at the bottom. **B.** On the top, secondary structures of NRU#10a and NRU#10b. On the bottom the sequence of NRU#10a and NRU#10b and the PhyloP score (based on the 100-way vertebrate alignment). Known structural and sequence motifs are marked in the sequence and in the structure.

Finally, we assessed whether the structure of the newly discovered NRUs was similar to the previously characterized ones, but aligning them to the previously mentioned alignment of the eight structurally conserved NRUs (NRU#1, NRU#3-8 and NRU#10). We were able to obtain a new consensus structure with the same short stem-loop structure but the consensus large stem-loop was smaller compared with the consensus structure without the new NRUs [Supplementary Figure S5A], suggesting that part of the consensus structure is not present in the newly identified NRUs. However, the structural similarity to the canonical consensus seems larger than that of NRU#2 and NRU#9. Based on these results, we conclude that the two newly identified NRU correspond to bona-fide functional elements in NORAD, and should be considered in future research.

## Discussion

NORAD is a critical regulator of genome stability. Moreover, human NORAD is dysregulated in various types of cancer and could be an excellent diagnostic or therapeutic biomarker in tumour cells [26]. Since lncRNAs have no protein-coding potential, their structure may be key for exploring their functionality. Despite this, there have been no studies of the *in vitro* structurome of NORAD. To fill this important gap, we performed nextPARS experiments that allowed us to study heat- or context-induced structural changes in NORAD at a single-nucleotide resolution. Moreover, we experimentally determined the *in vitro* secondary structure of NORAD and defined the core structural motifs conserved across NRUs. We identified two previously uncharacterized NRUs within an unusually long linker region. The addition of the two newly discovered NRUs results in all core units distributed along the whole NORAD sequence and placed at very regular distances. A possible explanation for this could be that the hairpin elements in the NRUs are interacting with each other or that they help to position the protein-binding sites at an advantageous distance from each other. We noticed that the NRU#10a was less conserved in their sequence compared to NRU10#b and the rest of the NRUs and, in contrast to the case of NRU#10b, we could not identify a putative SAM68-binding site sequence near the NRU#10a core.

The repetitive and modular nature of NORAD sequence allowed us to study several fragments at once and simplify the task of studying the structure of this long RNA transcript in its entirety. Our results indicate a stabilizing role of spacer sequences between NRUs, with their effect on the structure being larger than sharp shifts in temperature. How spacers stabilize NRUs is unclear. One possibility is that they favor interactions with other NRUs by placing them at some optimal distance, which would explain the regularity in spacer lengths. Alternatively, they may interact with some parts of the NRU structure, providing some additional stability. An RNA could fold in different structures inside the cell and many conformations for the same RNA coexist in equilibrium. Both for isolated NRUs and larger NORAD fragments experiments we observed that many sites exhibit clear signals for both single- and double-strand specific enzymes (nextPARS score near zero) which may result from noise, but also from the actual co-existence of alternative secondary structures during the experiment. Taking this into account, we estimated and compared the structural conformations of two NRUs and we detected structure motifs that were omitted when using the more stable structure. We revealed that these motifs were similar to the ones detected in some of the known NRUs. Moreover, we showed the importance of considering the ensemble of structures to compare structural elements in NRUs. Furthermore, a similar approach could be used to look for shared structural elements between different species to study the evolution of NORAD.

NORAD contains several SAM68-binding sites (UAAA motif) and the interaction between NORAD and the RNA-binding protein SAM68 is required for NORAD function. We compared the nextPARS data at different temperatures to evaluate the stability of SAM68-binding sites motif secondary structure. We revealed that positions of UAAA motifs were more robust to thermal shifts than other positions in the NRUs sequence. Interestingly, we observed a correlation between conservation and flexibility in UAAA motifs but only when we use nextPARS data from larger NRAD fragments experiments and not when using data from NRU-specific fragments.

For certain small RNAs, the *in vitro* folding landscape can recapitulate effectively the *in vivo* one, while for long RNAs the structures sometimes differ *in vitro* versus *in vivo* [47,48], These differences are mainly due to interactions with other molecules. *In vivo* structure probing may provide more accurate information on biologically relevant RNA structures [49,50]. In this work, we use only information from *in vitro* experiments, but we think that one future perspective could be the combination of these two experimental methods for obtaining the full picture of the RNA structurome. Nonetheless, both *in vitro* and *in vivo* approaches have limitations [51]. Furthermore, NORAD has sponge potential on several miRNAs and some of them were validated in different studies. Another future perspective could be to study the secondary structure of miRNA binding sites. We hypothesize that NORAD structure could modulate miRNA cleavage by facilitating or preventing accessibility of the miRNA to its binding site.

Taken together, our analyses contribute important novel insights into the structural organization of NORAD. Future studies should focus on determining how the different structural elements of NORAD interact with each other and with other molecules, and how different mutations may affect these interactions and the functions mediated by them. Only by understanding the complex relationships between, sequence, structure, and function will we be able to comprehend the roles of NORAD in the normal functioning of the cell and in diseases such as cancer. Eventually, this may open the door to design novel therapeutic strategies in the fight against cancer.

## Acknowledgements

UC was funded in part through H2020 Marie Skłodowska-Curie Actions (H2020-MSCA-IF-2017-793699) and MICINN (IJC2019-039402-1). TG group acknowledges support from the Spanish Ministry of Science and Innovation for grant PGC2018-099921-B-100, cofounded by European Regional Development Fund (ERDF); from the Catalan Research Agency (AGAUR) SGR423; from the European Union’s Horizon 2020 research and innovation programme (ERC-2016-724173); from the Gordon and Betty Moore Foundation (Grant GBMF9742) and from the Instituto de Salud Carlos III (INB Grant PT17/0009/0023 - ISCIII-SGEFI/ERDF).

