## Supplementary Figures for "Structural characterization of NORAD reveals a stabilizing role of spacers and two new repeat units"

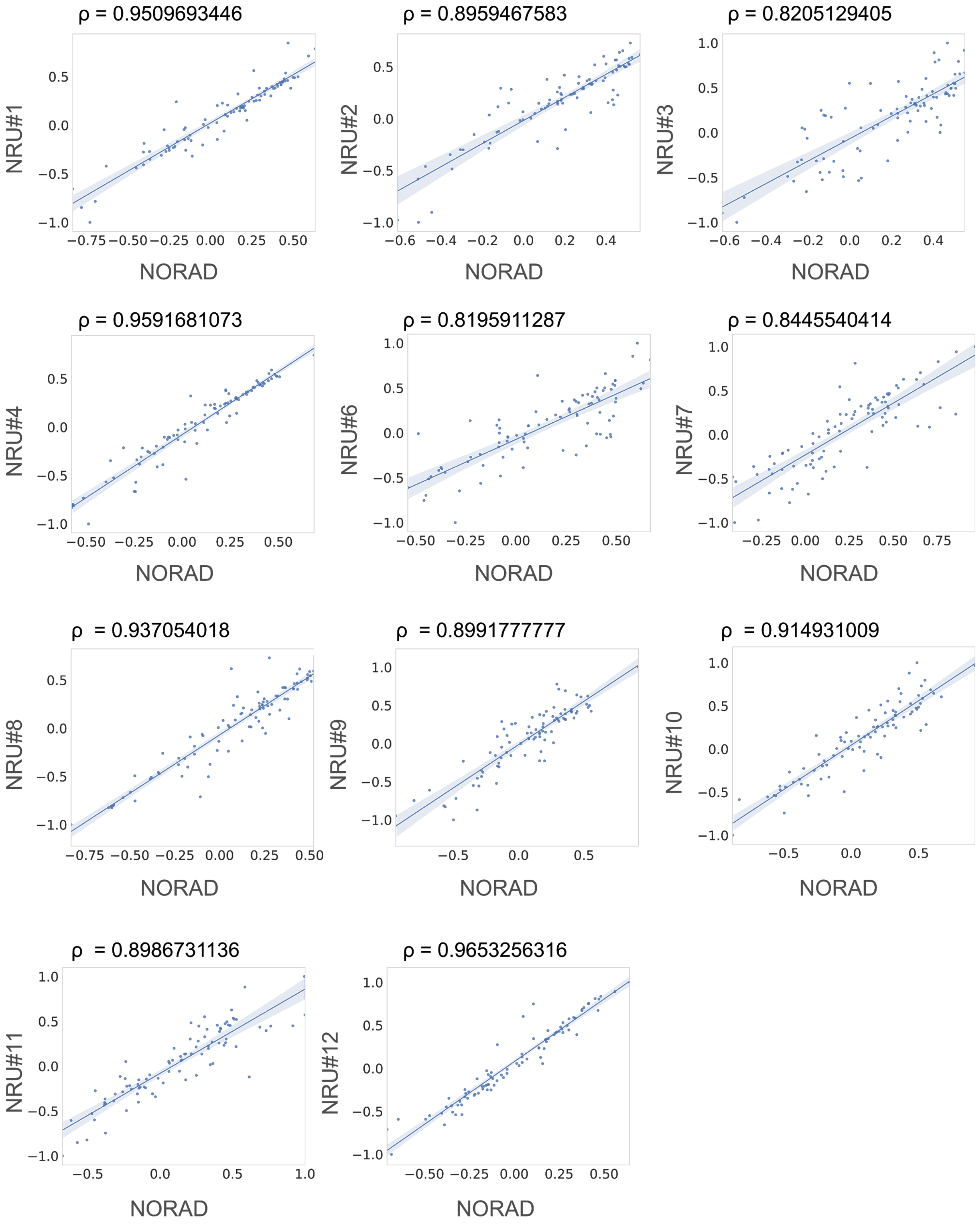


**Figure S1.** Pearson's correlation coefficient between nextPARS scores of NRUs within small or larger NORAD fragments.

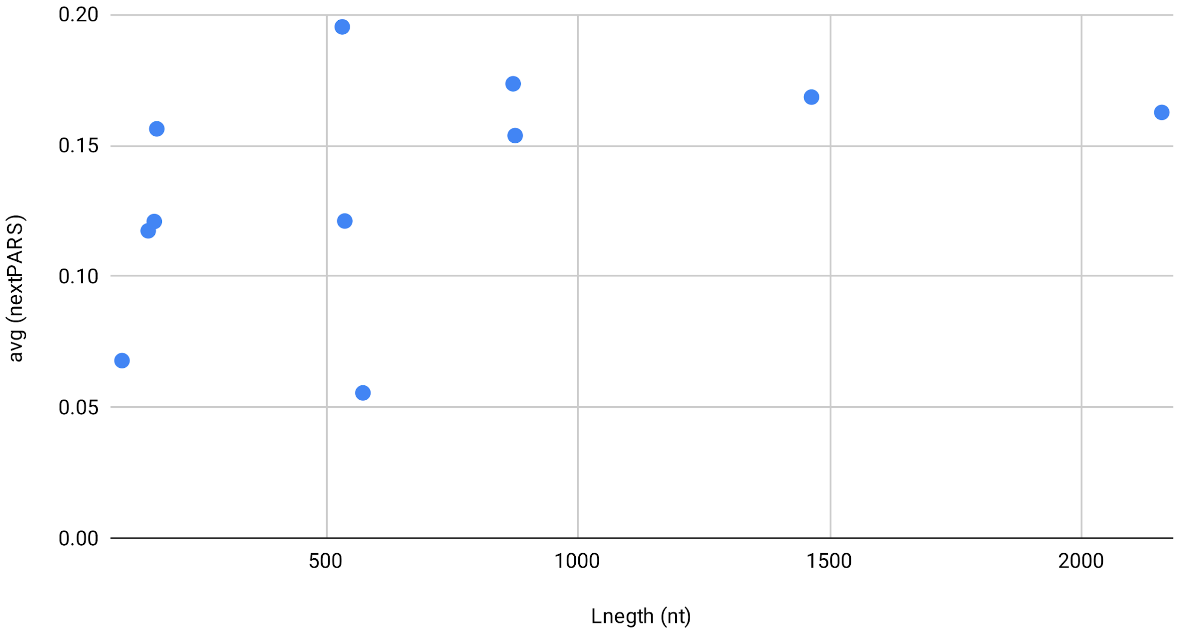


**Figure S2.**

Structural stability at different temperatures (average of nextPARS score) for previously performed nextPARS experiments of 11 RNAs molecules with variable size. Our results did not detect any statistically significant correlation between stability and sequence length, Pearson correlation coefficient 0.4428 (p-value 0.1725510616)

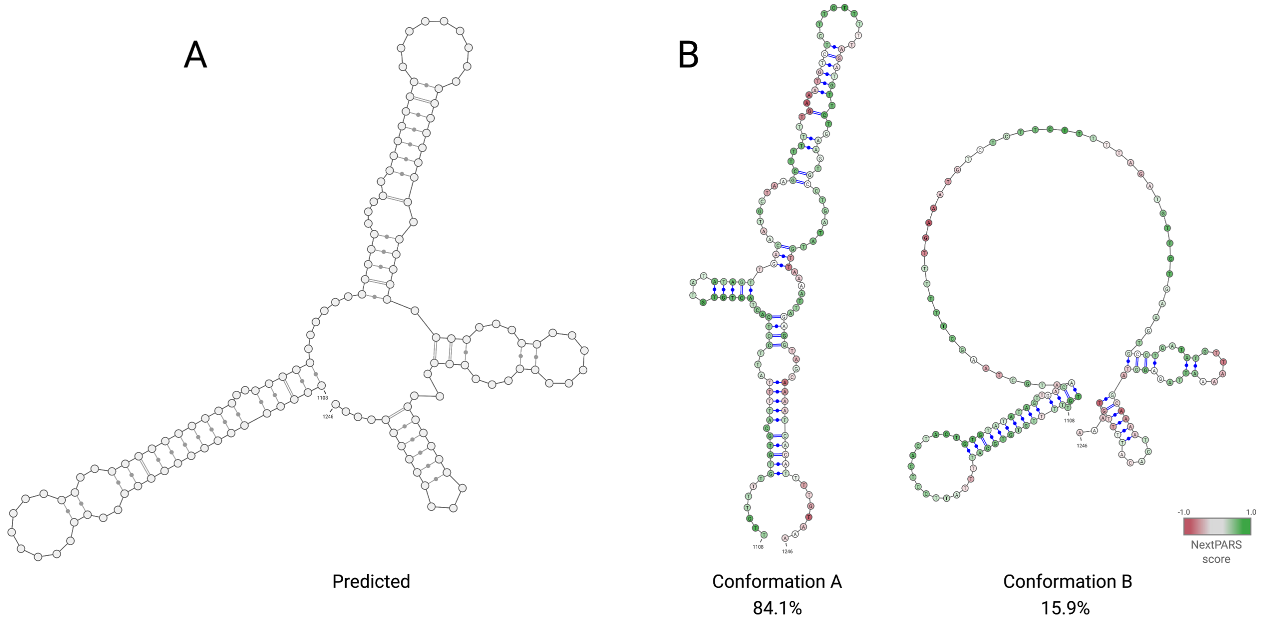


**Figure S3.** The Secondary structure of NRU#2 as predicted with RNAfold (**A**) in comparison with the two conformations predicted using Rsample combined with nextPARS data (B).

A

B

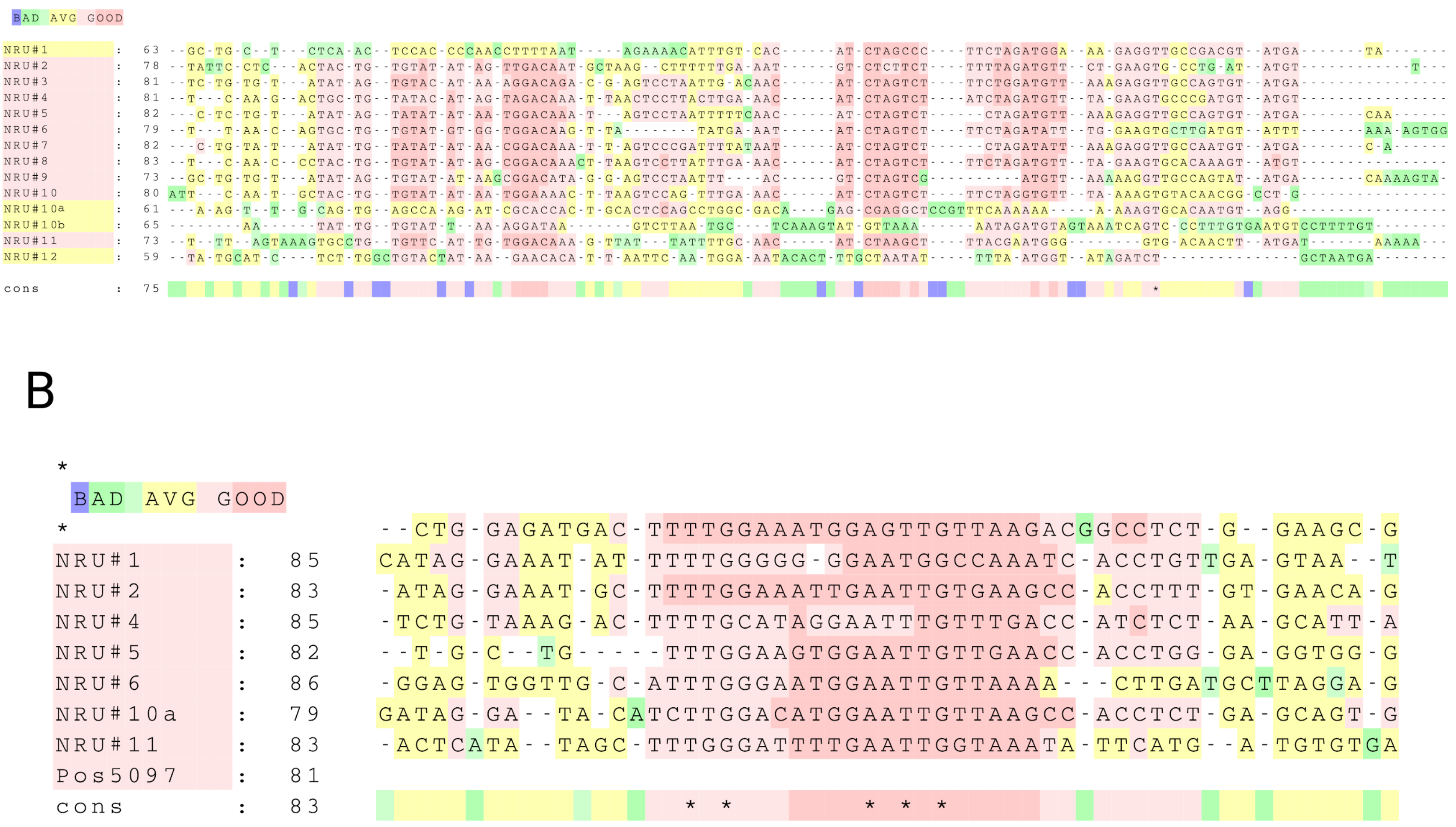


**Figure S4.** Multiple sequence alignments **(A)** of the core of the NRUs and **(B)** of a motif located at ~50 nt downstream of the end of some NRUs.

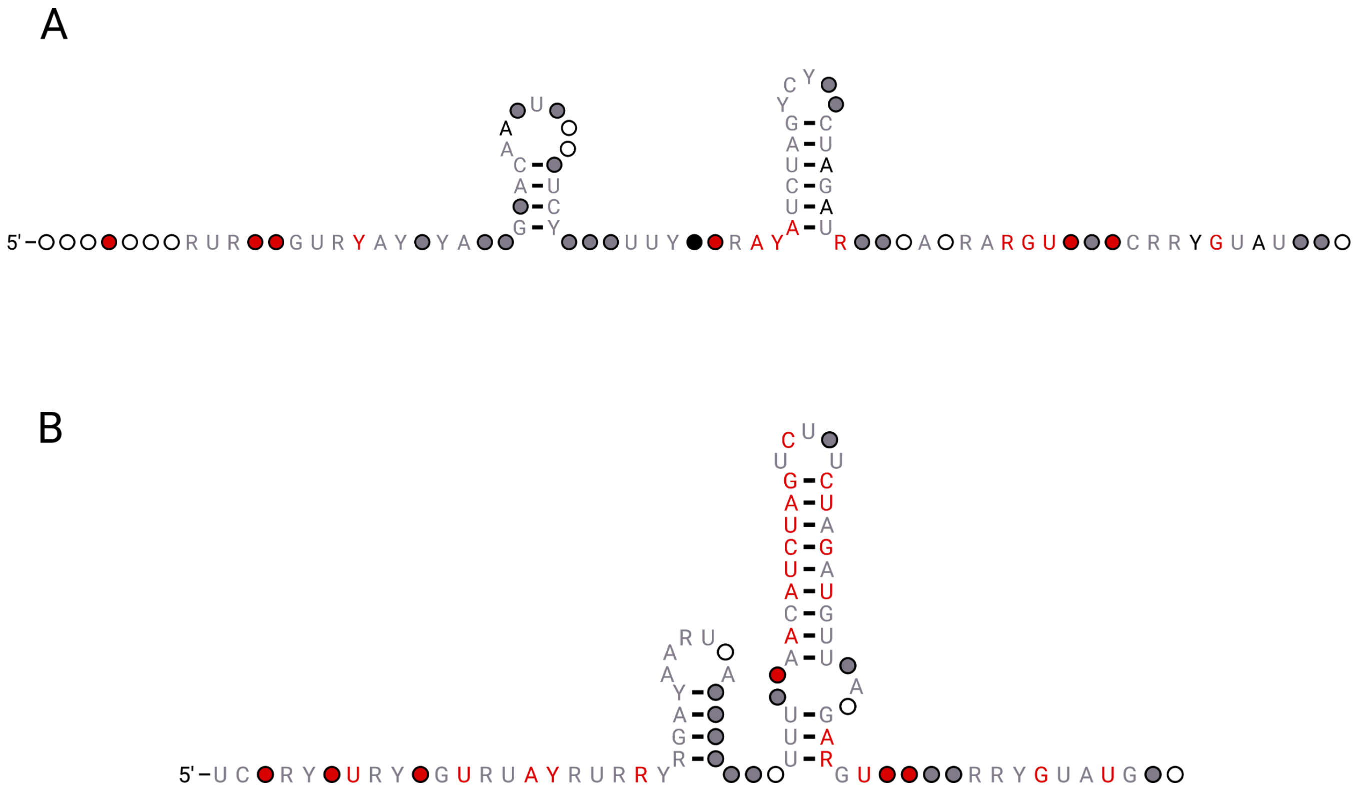


**Figure S5. (A)** The consensus secondary structure ofNRUs cores, derived by using a multiple MSA alignment to derive a from the eight NRUs that have a clear conserved long stem-loop hairpin (NRU#1, NRU#3-8 and NRU#10) with the addition of the two newly identified NRUs (NRU#10a and NRU#10b). **(B)** The same consensus secondary structure from the core of the NRUs, without the two newly identified NRUs.
