## Supplementary Table1 for "Structural characterization of NORAD reveals a stabilizing role of spacers and two new repeat units"

| **Fragment** | **Length (bp)** | **Forward primer (5’→ 3’)** | **Reverse primer (5’→ 3’)** |
| --- | --- | --- | --- |
| NORAD#1 | 1903 | CTCAAGTAATACGACTCACTATAGGGAGTTCCGGTCCGGCAG | CTATACTGTTCACAAAGGTGG |
| NORAD#2 | 1862 | CTCAAGTAATACGACTCACTATAGGGCCACCTTTGTGAACAGTATAG | GTCCTGATCTCTTGACCTC |
| NORAD#3 | 1614 | CTCAAGTAATACGACTCACTATAGGGGAGGTCAAGAGATCAGGAC | TACAGGCTTTTAAAATAACCTTTATTTTTTAAAAG |
| NRU#1 | 179 | CTCAAGTAATACGACTCACTATAGGGATTTACTGGCCGTTTATG | AAACAAATGAGAATTTACAAGATGTG |
| NRU#2 | 179 | CTCAAGTAATACGACTCACTATAGGGTTTAGAAGACATTTTCATATTC | AACAAAAAGGTATTTACAAAATGTGATTTTG |
| NRU#3 | 179 | CTCAAGTAATACGACTCACTATAGGGGCGAGGCAAGATGAATC | AAAACAAAATGTACAAAATATATTAGTTTAC |
| NRU#4 | 179 | CTCAAGTAATACGACTCACTATAGGGAAGACTCTTTCAAATTATAAC | AGCAAAAAGATATTTACAAAGTGGTATTTTAC |
| NRU#6 | 179 | CTCAAGTAATACGACTCACTATAGGGAAGATACCTCAGATGATA | TTTCCCATCAGTTTTTAAAAGCTATTTACAA |
| NRU#7 | 179 | CTCAAGTAATACGACTCACTATAGGGATCCATTGTGTATATGTTA | TTTTAACAAAGTGTACAAAATGTGTTAGTTT |
| NRU#8 | 179 | CTCAAGTAATACGACTCACTATAGGGGAAGACACTATCAGATATA | CAAAAGGATAGCTACAAAATGTGTTATTTAC |
| NRU#9 | 179 | CTCAAGTAATACGACTCACTATAGGGGGCACATTGACCATTGTCC | TGAATTTTAACACAAAGTGTACTCAATGTAG |
| NRU#10 | 179 | CTCAAGTAATACGACTCACTATAGGGAGAAGACACTCAAATTACA | CTCGGCCTCCCAAAGTGCTGGGATTACAGGC |
| NRU#11 | 179 | CTCAAGTAATACGACTCACTATAGGGCTAGACGATGGTTTTAGAT | ATTATAACAAAGGTATTTACAAATAGGCTAA |
| NRU#12 | 179 | CTCAAGTAATACGACTCACTATAGGGATGTGTCCATATATGTCCA | AAAATGAAACACACAGCAACAGAATACAGTA |

**Supplementary Table S1. PCR primers for NORAD fragments.** Underlined sequences from forward primers correspond to T7 promoter plus some additional bases to perform the *in vitro* transcription in a further step. The length of the amplicons shown in the table corresponds to the amplified region of interest without considering the extra-bases from T7 promoter.
