## Supplementary Table2 for "Structural characterization of NORAD reveals a stabilizing role of spacers and two new repeat units"

| **Fragment** | **Annealing temperature (ºC)** | | **Extension time (sec)** |
| --- | --- | --- | --- |
|  | **15 cycles** | **20 cycles** |  |
| NORAD#1 | 61.7 | 54.7 | 200 |
| NORAD#2 | 62.7 | 55.7 | 180 |
| NORAD#3 | 63.8 | 56.8 | 170 |
| NRU#1 | 60.7 | 53.7 | 20 |
| NRU#2 | 62.6 | 55.6 | 20 |
| NRU#3 | 60.0 | 53 | 20 |
| NRU#4 | 64.2 | 57.2 | 20 |
| NRU#6 | 63.9 | 56.9 | 20 |
| NRU#7 | 62.6 | 55.6 | 20 |
| NRU#8 | 65.3 | 58.3 | 20 |
| NRU#9 | 65.3 | 58.3 | 20 |
| NRU#10 | 74.7 | 67.7 | 20 |
| NRU#11 | 62.6 | 55.6 | 20 |
| NRU#12 | 66.6 | 59.6 | 20 |

**Supplementary Table S2. PCR conditions per each of the NORAD fragments amplified.** We performed touch-down PCRs for all fragments, reducing 0.5ºC of the annealing temperature in each cycle during the first 15 cycles. Except for the annealing temperatures and the extension time which are specified in the table per each of the fragments, the PCR conditions were the same for all fragments: 2 min at 94ºC; 15 cycles of 30 sec at 94ºC, 30 sec at each corresponding annealing temperature gradually reduced 0.5ºC in each cycle, the corresponding extension time at 72ºC; 20 cycles of 30 sec at 94ºC, 30 sec at each corresponding annealing temperature, the corresponding extension time at 72ºC; and a final extension of 5 min at 72ºC.
