## Supplementary Table3 for "Structural characterization of NORAD reveals a stabilizing role of spacers and two new repeat units"

**Supplementary Table S3. PCR primers for spike-in RNA fragments.** Underlined sequences from forward primers correspond to T7 promoter plus some additional bases to perform the *in vitro* transcription in a further step. The length of the amplicons shown in the table corresponds to the amplified region of interest without considering the extra-bases from T7 promoter.

| **Fragment** | **Length (bp)** | **Forward primer (5’→ 3’)** | **Reverse primer (5’→ 3’)** |
| --- | --- | --- | --- |
| TETp4p6 | 159 | CCAAGTAATACGACTCACTATAggaattgcgggaaaggggtcaac | gaactgcatccatatcaacag |
| TETp9-9.1 | 95 | CCAAGTAATACGACTCACTATAggacctctccttaatgggagc | ccaaaactaatcaatatactttc |
| hSRA | 875 | CTCAAGTAATACGACTCACTATAGGGCGCTTGGCGGAGCTGTAC | CACCACCGCAGAGATGTT |
| mSRA | 871 | CTCAAGTAATACGACTCACTATAGGGGTGCGGAAGTGGAGATGGC | TCATCTTTCCTACCCACTTCCC |
| ROX2 | 573 | CTCAAGTAATACGACTCACTATAGGGTGTTGCGGCATTCGCGGC | TTATTTGGCAATTGTTAAGTTTC |
| GAS5 | 537 | CTCAAGTAATACGACTCACTATAGGGCAGTGTGGCTCTGGATAGCA | TTGTGCCATGAGACTCCATCA |
| HOTAIR_NCBIBI_RINN | 2158 | ctcaagtaatacgactcactataggggactcgcctgtgctctggagc | ttttttttttgaaaatgcatccagatattaatatatc |
| 372_BRAVEHEART | 532 | CTCAAGTAATACGACTCACTATAGGGGATGGAACAGGAGGAGCATC | TCAGCTAGGTAAACTAAAAGCCA |
| 373_BRAVEHEART | 1463 | CTCAAGTAATACGACTCACTATAGGGTCCAGTGCCAGTTGCTTAAA | GAATGGGGATAGGGCATTCT |
| B2 | 147 | CTCAAGTAATACGACTCACTATAGGGCCGACTGCTCTTCCGAGGTC | TTTAAAGATTTATTTATTTATTATATGTAAG |
| U1 | 164 | CCAAGTAATACGACTCACTATAGGGATACTTACCTGGCAGGGGAGATACC | CAGGGGAAAGCGCGAACGCAGTCCC |
